# How do we segment text? Two-stage chunking operation in reading

**DOI:** 10.1101/806190

**Authors:** Jinbiao Yang, Qing Cai, Xing Tian

## Abstract

*Chunking* in language comprehension is a process that segments continuous linguistic input into smaller chunks that are in reader’s mental lexicon. Effective chunking during reading facilitates disambiguation and enhances efficiency for comprehension. However, the mechanisms of chunking remain elusive, especially in reading given that information arrives simultaneously yet the written systems may not have explicit cues for labeling boundaries such as Chinese. What are the mechanisms of chunking operation that mediates the reading of the text that normally contains hierarchical information? We investigated this question by manipulating the lexical status of the chunks at distinct levels of grain-size in four-character Chinese strings, including the two-character local chunk and four-character global chunk. Participants were asked to make lexical decision on these strings in a behavioral experiment, followed by a passive reading task when their electroencephalography (EEG) were recorded. The behavioral results showed that the lexical decision time of lexicalized two-character local chunks was influenced by the lexical status of four-character global chunk, but not vice versa, which indicated that the processing of global chunks possessed priority over the local chunks. The EEG results revealed that familiar lexical chunks were detected simultaneously at both levels and further processed in a different temporal order -- the onset of lexical access for the global chunks was earlier than that of local chunks. These consistent behavioral and EEG results suggest that chunking in reading occurs at multiple levels via a two-stage operation -- simultaneous detection and global-first recognition.

**Significance Statement:** The learners of a new language often read word by word. But why can proficient readers read multiple words at a time? The current study investigates how we efficiently segment a complicate text into smaller pieces and how we process these pieces. Participants read Chinese strings with different structures while their key-press responses and brain EEG signals were recorded. We found that texts were quickly (about 100 ms from their occurrences) segmented to varied sizes of pieces, and larger pieces were then processed earlier than small pieces. Our results suggest that readers can use existing knowledge to efficiently segment and process written information.

## Introduction

Reading is arguably one of the unique human intelligences. Yet how we process written texts remains elusive. For instance, how can we comprehend a complex sentence? A sentence consists of many letters/characters that form a hierarchical structure of text chunks (e.g. morphemes, words, and phrases). Readers need to incrementally segment a complex sentence into smaller chunks that map onto their mental lexicon. This process is termed as text *chunking* (Gobet, Lloyd-Kelly, & Lane, 2016; Reali & Christiansen, 2007). What are the small chunks during chunking? How do we process the chunks? To answer those questions, this study investigated the cognitive procedure of text chunking.

Words and their sub-level (morphemes) are usually assumed as the basic units in reading models in psycholinguistics and computer science (Coltheart, Rastle, Perry, Langdon, & Ziegler, 2001; McClelland & Rumelhart, 1981b; Taft, 2013). However, eye-tracking studies suggested we can perceive the text information longer than a word at one time (Rayner, 1998). Our working memory also allows us to remember familiar multiple words (Miller, 1956). Even more, multi-word expressions can be stored in our mental lexicons (Arnon & Snider, 2010/1; Siyanova-Chanturia, Conklin, Caffarra, Kaan, & van Heuven, 2017). These studies suggest the multi-word representations and the beyond-word processing are feasible. Moreover, relying on larger chunks effectively reduces the cognitive load while processing sentences: fewer chunks to be interpreted and integrated (Philippe Blache & Rauzy, 2012; Nick C. Ellis, 2003; Krishnamurthy, 2003). Furthermore, the semantic combination of constituents can be different from the holistic meaning (Goldberg, 1995). One extreme example is idioms, as the metaphors of an idiom can be distinct from their literal meanings of smaller constituents. Multi-word representation is required in certain contexts to avoid ambiguity. Therefore, multi-word chunks, as well as word chunks, could be the units during chunking.

How do we integrate the processes on word and multi-word chunks that co-exist during chunking? The studies of compound words (a single lexical entity but consists of more than one root morphemes; e.g. “flagship”) may offer hints. According to the dual-route models of compound-word processing, both the whole word and its constituents are processed at the same time or are selected to process each level flexibly (Andrews, Miller, & Rayner, 2004; P. Blache, 2015; Koester, Gunter, & Wagner, 2007; MacGregor & Shtyrov, 2013; Semenza & Luzzatti, 2014). In a similar vein, we hypothesized that all the familiar lexical chunks, no matter which level it is, could be processed simultaneously. More specifically, the detection of chunks would be the first step in chunking, and the detection of chunks at multiple levels would occur at the same time, as the early lexical familiarity checking assumed in the E-Z reader model (Reichle, Rayner, & Pollatsek, 2003).

How does the multi-level operation unfold in the chunking process? Which level has the priority after being detected? The word superiority effect indicates that the recognition of letters within words is better than letters in nonwords or stand-alone letters (Reicher, 1969). It suggests that the word has priority over the letter in reading. Similarly, the processing priority of global chunks can reduce the steps of integration and avoid the ambiguity to enhance the efficiency of language processing (Philippe Blache & Rauzy, 2012; Nick C. Ellis, 2003; Krishnamurthy, 2003). Generalizing from the word superiority effect, we hypothesized that global chunks take the priority over the parts and would be initiated first in the processing stage after detection.

In this study, we used Chinese four-character strings to investigate the chunking operation in reading. Chinese written system is a good model for observing multi-level chunking because Chinese does not have explicit word boundaries. Each Chinese character is a basic lexical unit with similar length and four characters can form two levels of chunks -- chunks with 2 characters (hereafter as the *local level chunks*) and a chunk with 4 characters (hereafter as the *global level chunk*). The lexicality was manipulated at both levels so that four types of stimuli were included (phrase, idiom, random words, and random characters). In the behavioral experiment, we investigated the interaction between the global and local chunks in reading by a lexical decision task at different levels of chunks. Moreover, an EEG experiment was carried out to investigate the temporal dynamics of detection and recognition stages in the multi-level chunking operation.

## Methods

### Participants

Twenty-one healthy native Chinese speakers (10 males, mean age 21 years, range 18-30 years) with normal or corrected-normal vision participated in both behavior and EEG experiments for financial compensation. The experiments were approved by the Research Ethics Committee of [Author University]. Written informed consents were obtained for all participants before the experiments.

### Stimuli

All stimuli are four-Chinese-character strings. Two factors are included when designing these stimuli. The first factor is the *chunk size* that contains two levels -- a global size of 4 characters and a local size of 2 characters. The second factor is *lexicality* (word or nonword) at each chunk size. These two factors are fully crossed and yield 4 types of stimuli. We denote *chunk size* using upper case letters -- ‘G’ for global and ‘L’ for local, and use lower case letters for lexicality in each chunk size (‘w’ for word and ‘n’ for nonword). For example, ‘GnLw’ stands for the condition of stimuli that are four-character nonword at the global level made of two two-character words at the local level. The four types of stimuli are listed in Table 1.

**Table 1.**
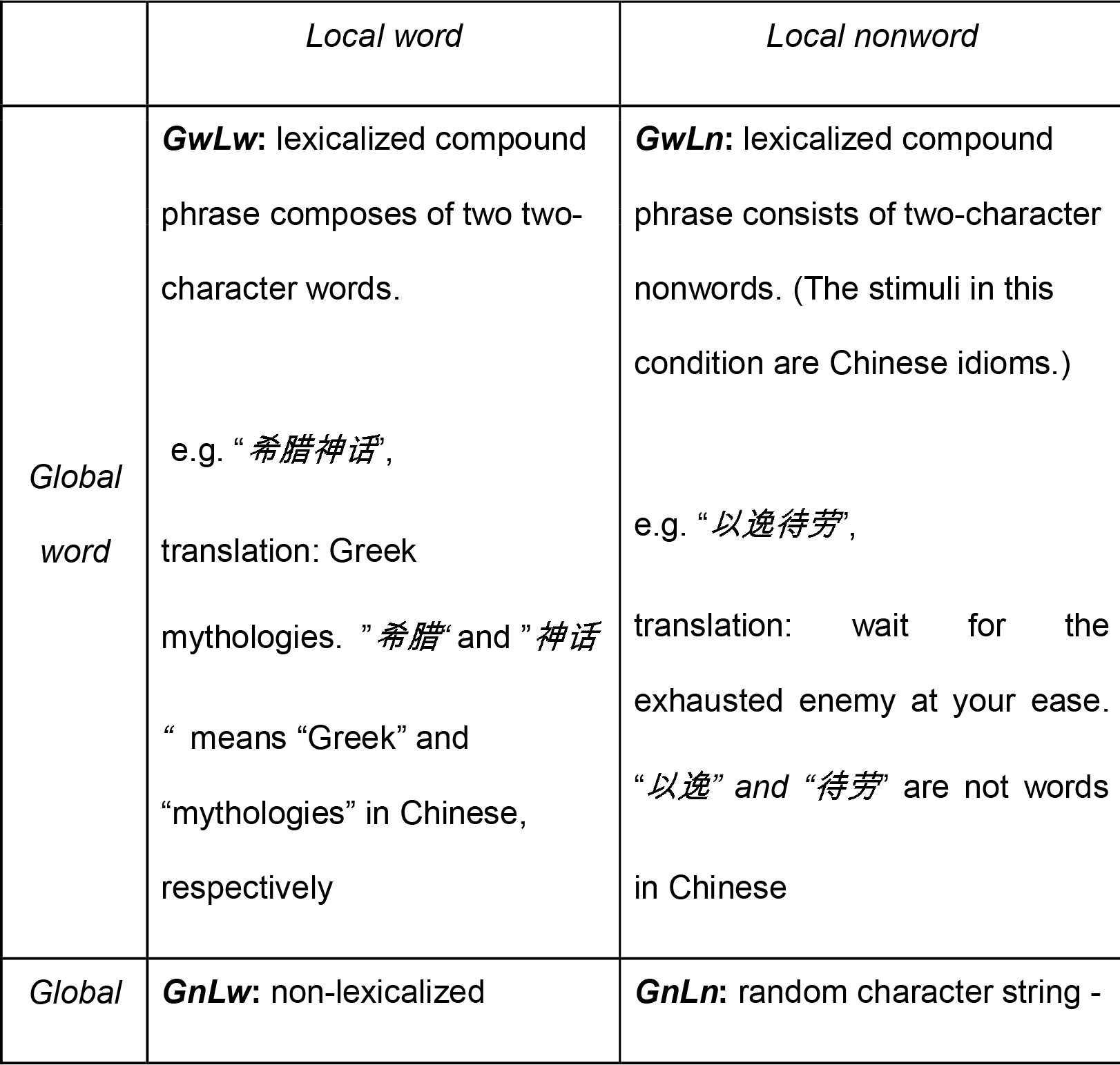
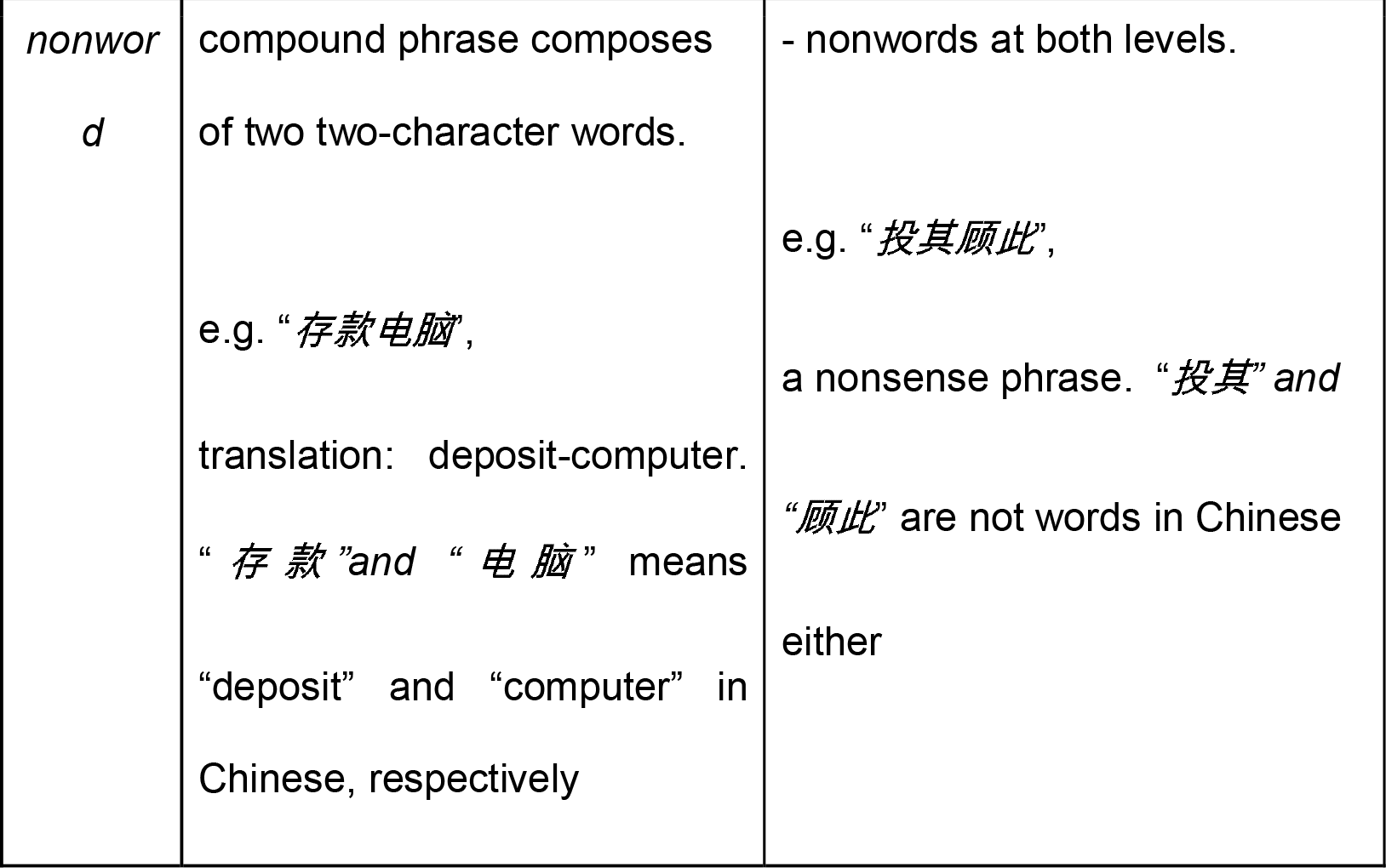
stimuli description.

We selected and created all stimuli with following steps. We extracted the *GwLw* and *GwLn* stimuli from a database of Sogou Pinyin (a popular product of Chinese character input method) and a database of Chinese characters (CharDB: Data version: 0.98.1; Program version: 0.97.2; http://chardb.science.ru.nl/). All the *GwLw* and *GwLn* stimuli satisfied the following criteria at the global level:1) noun^1^; 2) high-frequency^2^; and 3) no duplicative characters (e.g., “*高高兴兴*”, translation: happy). In addition, the *GwLw* stimuli satisfied the following criteria at the local level: 1) both two-character words were nouns, and 2) high-frequency words. Moreover, the lexicality of *GwLn* stimuli at the local level were verified by checking the first two or last two characters’ combination did not exist in the Sogou Pinyin database. These selection criteria made the *GwLw* and *GwLn* stimuli consistent in all aspects except the lexical status at the local level.

The GnLw stimuli were created by randomly pairing two different high frequency two-character words. So the only difference between the *GwLw* and *GnLw* was the lexical status at the global level. Finally, the *GnLn* stimuli were created by randomly mixing four different characters, and none of the first or last two characters’ combination existed in the Sogou Pinyin database. Characters used in all stimuli have log frequency^3^ ranging from 3.011 to 5.344, with stroke counts ranging from 4 to 13.

The distinction between word and nonword were further controlled by familiarity. Twelve participants who were not in the main experiment were asked to rate the familiarities of either the entire four-character or the constituents of two-character strings as being words or not. The rating range was from 1 to 5, where 1 stands for unfamiliar strings/nonwords and 5 for familiar words. The strings that were rated from 2 to 4 were removed and remained the stimuli that were either very familiar words or very unfamiliar nonwords in a pool. Eighty stimuli in each condition was randomly selected from the pool and used in this study.

### Procedure behavioral experiment

In each trial, participants were first asked to focus on a cross presented at the center of the screen. After 400 ms the fixation cross disappeared, and a 4- character string was shown until response. A line was also appeared either under the entire 4-character string or under the 2-character string (the first or the last 2 characters). Participants were asked to make a lexical decision about the underlined string, either the entire string (*global task* henceforth) or the constitute of first or last 2-character string (*local task* henceforth) by pressing either “F” or “J” on the keyboard as fast as possible. Participants had maximum 3 seconds to respond. Responses and reaction time were collected. Response keys were counterbalanced across participants. The inter-trial intervals were randomly selected from a range from 800 to 1000 ms.

Four stimuli types (*GwLw*, *GwLn*, *GnLw*, *GnLn*) were fully crossed with task types (global task vs. local task) and yield 8 conditions. 320 trials were included in this experiment. Half of trials were randomly selected and used in the global task and the other half in the local task. The order of conditions was randomized. The experimental presentation was programmed on a Python package – Expy (https://github.com/ray306/expy), which is a software for presenting and controlling psychological experiments.

#### Behavioral data analysis

All participants had response accuracy exceeding 85%, and the average of accuracy was 92%. No participant’s data was excluded. Trial with error responses were removed before analysis. We applied a repeated measures three-way ANOVA on the reaction time data with factors of *global-level lexicality, local-level lexicality*, and *task*, followed by planned t-tests for testing specific hypotheses.

### Procedure EEG experiment

The same group of subjects participated in the EEG experiment. The EEG experiment shared the same stimuli list with the behavioral experiment, but both the procedure and the task are different. First, the display of each character string lasted for 300ms. Participants were asked to read the underlined parts of the stimuli (in order to keep their attention on the stimuli), but they did not perform any lexical decision task. We used all 320 strings with 80 for each stimuli type in the *global task* and repeated once in the *local task*. Moreover, 320 four-symbol strings were included as the visual baseline in the EEG experiment. The symbols in a symbol string trial were randomly sampled with replacement from four symbols (“□”, “△”, “⍰”, and “○”). Underlines were included in the symbol trials similar as in the global and local tasks in experimental trials. Half of trials were randomly selected and used in global task and other half in local task. To guarantee participants’ attention on the stimuli, we randomly inserted strings of digits for 100 ms and participants were asked to report the underlined digits by pressing number buttons on a keyboard. About forty-eight number-report trials were presented to each participant.

#### EEG recording

EEG signals were recorded with a 32-electrode active electrodes system (actiChamp system, Brain Products GmbH, Germany). FP1 and FP2 were used to monitor vertical eye movements. Electrode impedances were kept below 10 kΩ. Data were continuously recorded in single DC mode. Data were sampled at 500 Hz, online referenced to the Cz.

#### EEG data analyses

EEG data were preprocessed using EEGLAB (Version 13.5.4b; Delorme & Makeig, 2004). Data were band-pass filtered (0.1–30 Hz, Hamming windowed sinc FIR filter), and re-referenced to the average reference. The preprocessed data were epoched between −200ms and 800ms relative to the onset of strings, and baseline-corrected using the 200ms pre-stimulus period. The trials with eye blinks were rejected if the amplitude within the 1000ms epoch exceeded ±50μV, then the remain trials with apparent noise were rejected manually. About 15% of trials were rejected. Five participants who produced a large number of artifacts or showed continuous alpha waves were excluded from further analysis. Epochs in each condition were averaged and created an ERP response.

Our analysis focused on topographic patterns across all sensors rather than the response amplitude in selected groups of sensors, as it can provide more holistic and unbiased information. We used multivariate instead of univariate methods to test our hypotheses because multivariate methods can collectively reflect spatial and temporal information and offer more power to test psychological and neuroscience hypothesis by overcoming problems such as individual differences, sensor selection and reference selection in EEG (Murray, Brunet, & Michel, 2008; Tian & Huber, 2008; Tian, Poeppel, & Huber, 2011; Yang, Zhu, & Tian, 2018). Three multivariate based methods as follows were applied.

##### 1) Clustering

A clustering method on the ERP topographic responses were implemented. This unsupervised machine learning method groups data across all conditions by forming temporal clusters based on the similarity of their topographical patterns. Compared with TANOVA, this clustering method is a data-driving method, in which it explores the pattern similarity in topographies in all conditions. The clustering algorithm organizes data at different time points into distinct clusters, so that we can explore the temporal dynamics of pattern changes. Moreover, if one considers the clusters reflecting different processing stages, this analysis can identify the processing stages in each condition, and display the temporal differences of any specific stage among conditions. We used K-means, the most popular algorithm for clustering.

The procedure of clustering algorithm covers three steps: a) averaged EEG data across all participants to get ERPs at each time point for each group; b) defined ERPs at each time point in each group as a sample, and the amplitudes of thirty-two electrodes were used as features in a sample; c) K-means algorithm was carried out at all samples to get five clusters. The number of clusters was initially two, and increased until the result included more than one clusters at the baseline stage.

##### 2) Analysis of amplitude differences in topography

Topographies represent the response amplitude distribution across all sensors at a given time point. The changes of amplitude distribution in topographies can reflect the dynamics. The spatial extent of experimental effects can be estimated by the distribution and number of sensors that are significantly different between conditions. At a step size of 20 ms, we checked the topographies and the significant electrodes (Yang et al., 2018) to obtain the differences between conditions. Because there were thirty-two comparisons on each topography, the *p*-values were corrected by false discovery rate (FDR).

##### 3) Topographic Analysis of Variance

We further investigate the patterns of topographies by considering all sensors at the same time to infer the differences of underlying neural processes across conditions. A single index was calculated to indicate the topographic information. Mathematically, each topography can be viewed as a n-dimensional vector, where the n equals to the number of sensors. The divergence between the topographies of two experimental conditions can be quantified by the cosine value of the high-dimensional angle between two vectors (Tian & Huber, 2008). The cosine distance has a range from 0 to 2, where 0 stands for identical topographies and 2 exactly opposite patterns. Note that the cosine distance represents the similarity between the response patterns in topographies and is free from the difference of response magnitude because the measure of cosine distance is normalized by the vector length. To statistically test the cosine distance between topographies and to infer the underlying neural processing in different conditions, we applied an algorithm named Topographic Analysis of Variance (TANOVA) (Brunet, Murray, & Michel, 2011; Lange, Perret, & Laganaro, 2015; Murray et al., 2008; Tian & Huber, 2008; Tian et al., 2011). In TANOVA, null hypothesis distribution is generated by shuffling the condition labels, and we here shuffled the condition labels on the subjects’ ERPs using EasyEEG toolbox ((Yang et al., 2018), Strategy 3, shuffle times: 500, window size: 10 ms).

The test of TANOVA involved comparisons on multiple timebins, we corrected the multiple comparison by considering the temporal cluster size -- the number of adjacent significant timebins. Specifically, only 2 or more continuous timebins that reach the significant level of p<0.05 were considered as true significance.

## Results

### Behavioral experimental results

Reaction time was subject to a repeated measures three-way ANOVA with the factors of *global-level lexicality*, *local-level lexicality,* and *task*. The main effect of *global-level lexicality* was significant [*F*(1,19)=13.69, *p*<0.01], suggesting that participants took longer time to identify global-level words than global-level nonwords. The main effect of *local-level lexicality* is marginal significant [*F*(1,19)=3.815, *p*=0.066], suggesting the chunk at global level influenced lexical decision more than chunks at local level. However, the main effect of *task* is not significant [*F*(1,19)=0.389, *p*=0.54], suggesting different tasks that require participants to respond to either global or local chunks have similar level of difficulty. More importantly, all three two-way interactions are significant, *global-level lexicality* * *local-level lexicality* [*F*(1,19)=64.417, *p*<0.001], *global-level lexicality* * *task* [*F*(1,19)=27.782, *p*<0.001], and *local-level lexicality* * *task* [*F*(1,19)=92.913, *p*<0.001].

Planned post-hoc T-tests were further carried out in each factor to specify the observed significant interactions. First, we examined how the global information affect processing at the local level (Fig. 1A). In the local task, the reaction time in *GnLw* was significantly longer than that in *GwLw* (*t*(19)=5.7145, *p*<0.001, difference=55.3683), suggesting that the nonwords at the global level significantly slowed down the lexical decision of words at the local level. Moreover, the reaction time of *GwLn* was significantly longer than *GnLn* in the local task (*t*(19)=3.6214, *p*=0.002, difference=54.1903), suggesting that the words at the global level also slowed down the lexical decision of nonwords at the local level. That is, whenever there is a mismatch in lexical status across levels, global chunks affected lexical decision of local chunks. Second, we examined how the local chunk could affect processing at the global level (Fig. 1B). In the global task, we didn’t find significant difference between reaction time to *GwLw* and *GwLn* [*t*(19)=−1.0091, *p*=0.32], suggesting the lexical status of local chunks cannot affect lexical decision of words at the global chunks. The reaction time of *GnLw* was significantly longer than that of *GnLn* in the global task (*t*(19)=6.6874, *p*<0.001, difference=112.8455), suggesting the decisions of global chunks and local chunks could be parallel when global decision took too long, and the lexical information at the local level may leak through to the processing of global chunks and influence the decision of nonwords. We further test this parallel processing dynamics in EEG experiment.

**Figure 1.**
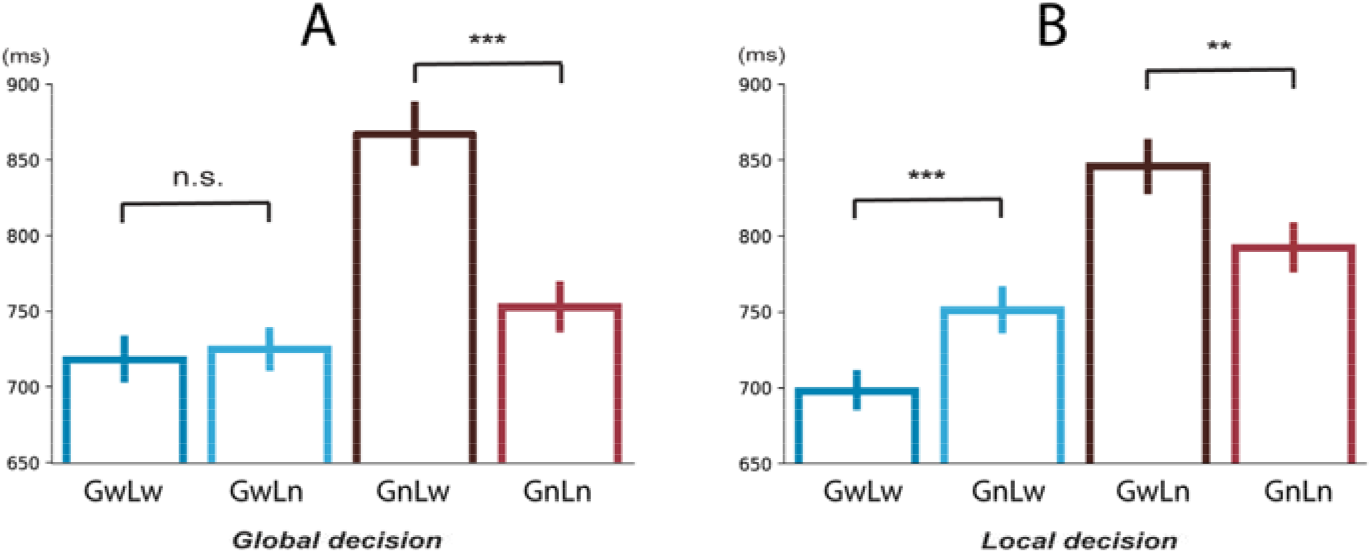
Results of behavioral experiment. A) and B) depict the reaction time results in the global and local tasks, respectively. In each plot, condition labels are provided along the x-axis. Error bars represent +/− one standard error of the mean (SEM). Each planned paired test was represented by the line linking two bars. n.s. stands for not significant, ** stands for p < 0.01, and *** stands for p < 0.001.

The reaction times were not different between the trials with underlines either below the first or the last two characters (*p*=0.44), suggesting the positions of stimuli that were relevant to task did not affect response speed.

### EEG results

#### Clustering revealed distinct stages of chunking

We first carried out the clustering analysis to explore the dynamics of ERP responses. The clustering algorithm aimed to separate the continuous ERP responses into distinct stages based on common features observed across time. As shown in Figure 2, the clustering results were reliable as similar clusters were observed continuously in each condition.

**Figure 2.**
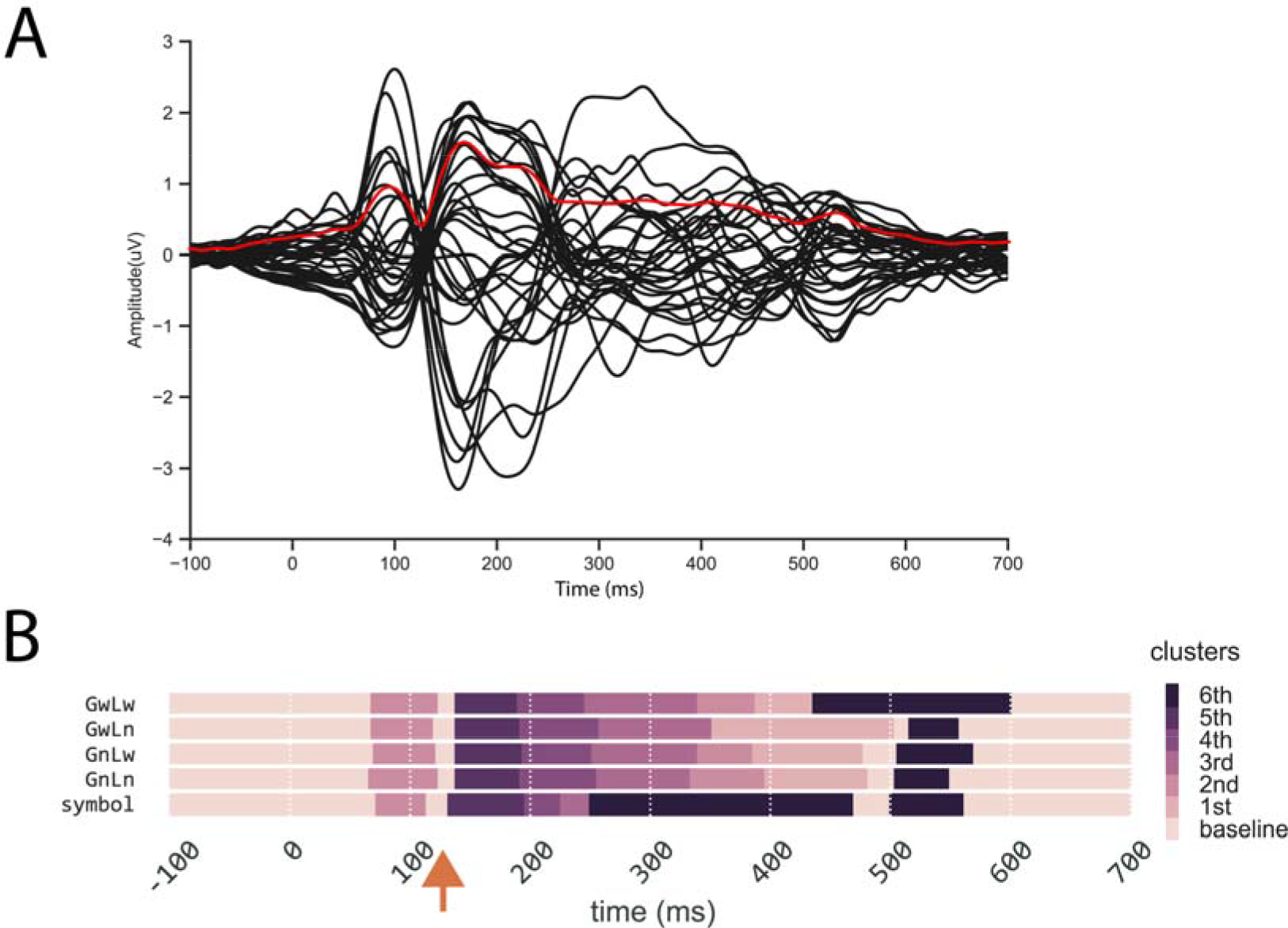
The dynamics of ERP responses and clustering results. (a) ERPs in each of thirty-two electrodes (black curves), root mean square (RMS, red curve). (b) Temporal clustering results of topographies for four conditions (GwLw, GwLn, GnLw, GnLn) and a baseline symbol condition (Symbol). Different colors represent distinct clusters. Samples in the same color but at different time points indicate that they are grouped into the same cluster -- sharing similar features but occurring at different times. The temperature of colors represents the rank of the cluster distance relative to the Cluster baseline (cluster defined by the baseline period). About 80 ms after stimulus onset, a novel cluster (Cluster 2nd) appears at the same time across 5 conditions, followed by another new cluster. However, in the symbol condition the Cluster 2nd appears earlier with much shorter duration than 4-character string conditions.

More importantly, clear temporal profiles were revealed by the clustering analysis in all conditions. First, the same cluster was observed in the baseline period till around 80 ms after stimulus onset, as well as the end of epochs (about 600 ms after onset) among all types of stimuli. The clustering in these periods was presumably because few cognitive processes that relate to the stimuli or task were available or manifested in the ERP topographies. Second, a novel cluster (Cluster *2nd*) appeared after 80 ms across 5 conditions. The clustering spanned in similar latencies as N1/P2 components, presumably reflecting visual processing. However, different dynamics was observed across conditions after 200 ms. The Cluster *3rd* appeared earlier in the *symbol* condition with much shorter duration than the four experimental string conditions in which the Cluster *3rd* appeared around 250 ms and lasted about 90 ms. Moreover, in the *symbo*l condition, the Cluster *3rd* was immediately followed by the Cluster *6th* that did not appear till 500 ms in the four experimental string conditions (except in *GwLw condition around 430ms*). The early start and long lasting Cluster *6th* in the *symbol* condition was accompanied with the missing of the Cluster *1st* and Cluster *2nd* that appeared around 320 ms and lasted till 450 ms in other conditions.

More interestingly, a 15 ms period around 130 ms was labeled as the Cluster *baseline* that was grouped in the baseline and the end of epoch periods. This formed a short gap that broke the early processing into two stages. We focused on the components in early timing to further investigate the underlying processes of chunking operation.

#### Chunk detection in the earliest stage

To test the hypothesis about the lexical detection in the earliest stage, we carried out two types of analyses to investigate the lexicality effects regardless at the global or local chunk levels. First we check the response amplitude differences among conditions. Compared with the responses to *GnLn* strings that contained no lexical chunks at either level, response amplitudes in conditions that include lexical chunks (*GwLw, GwLn, GnLw*) did not show any statistically significant differences in any sensors after multiple comparison correction (Fig. 3a). However, the difference topographies showed distinctive patterns of amplitude distribution (higher on left frontal area and lower on occipital area, highlighted in a red box in Fig. 3a). Therefore, we investigated the topographic patterns to infer the different configuration of neural processing across conditions.

**Figure 3.**
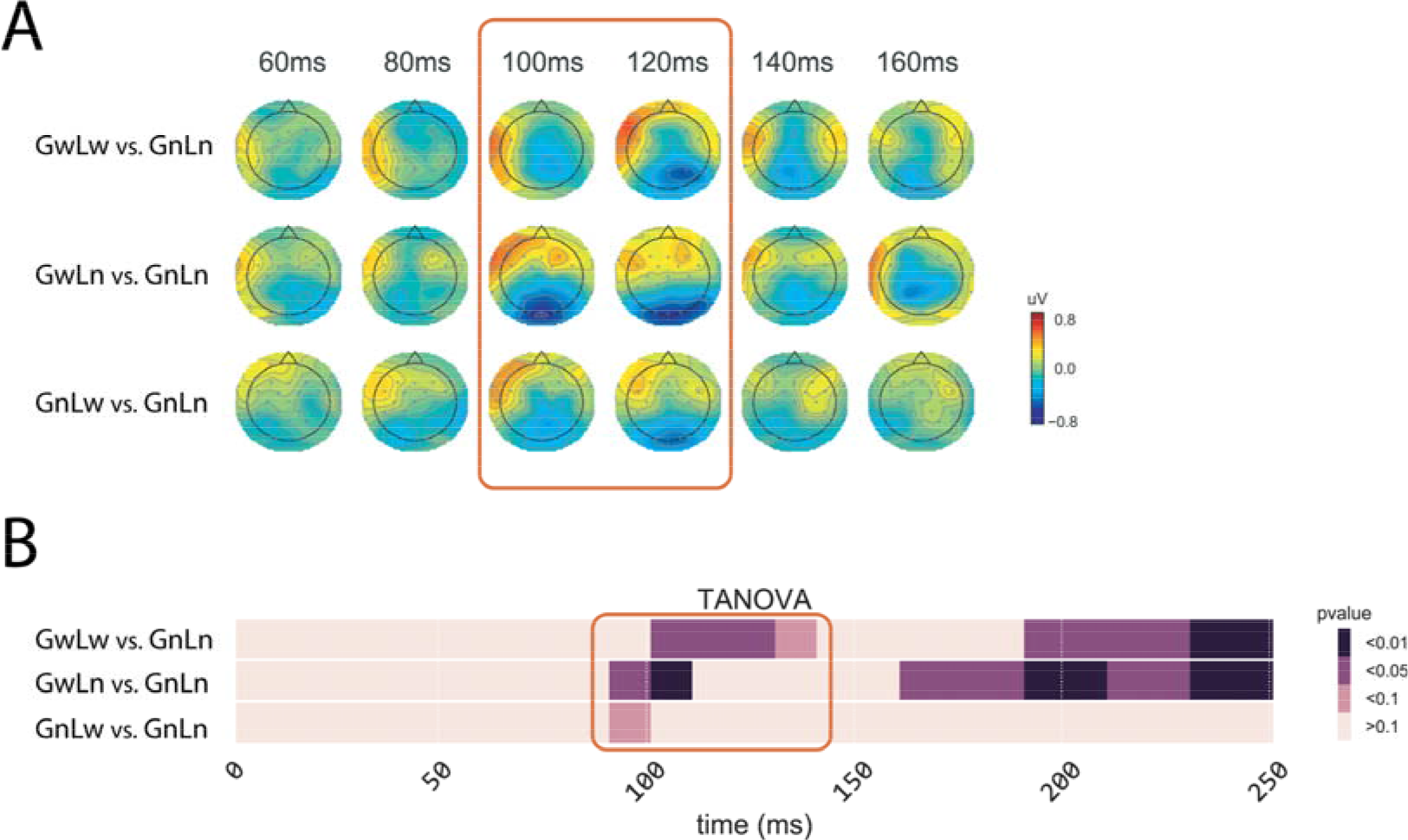
The effects of lexicalized chunks revealed in the paired comparisons between the GnLn condition and the other three lexical conditions (GwLw, GwLn, and GnLw). (a) Topographical comparisons of response amplitude. Each row shows a comparison across time. The color scheme depicts the differences in response amplitude between conditions. The red box highlights the earliest appearance of amplitude differences. No statistical significant difference was found on the electrodes after the multiple comparison correction (FDR). (b) The temporal dynamics of TANOVA results. The red boxes highlight the earliest latency when the significant differences were observed. The scale that includes p<0.1 was chosen only for visual inspection of weak effects in the GnLw condition.

The analysis of TANOVA revealed significant differences between the topographies of *GnLn* and responses patterns in *GwLw*, *GwLn* and *GnLw* conditions (Fig. 3b highlighted in the red box). The differences were first observed at 90 ms after stimulus onset. The differences were largest in the *GwLn* condition as the significant level at p<0.01 for the following 20 ms. The pattern differences were also observed in the *GwLw* condition, but the significant differences started later at 100 ms and lasted for 30 ms. The differences were weakest in the *GnLw* condition as they were only marginal significant from 90 to 100 ms. The TANOVA results suggested that the early stage process that before 130 ms related to the detection of lexicality as the differences were observed when lexicalized chunks at either level compared to non-lexicalized *GnLn* condition, and the global level information may facilitate the detection as the effect size ranked from biggest to smallest in the order of *GwLn*, *GwLw* and *GnLw*.

#### Processing of chunks at different levels

We also applied responses amplitude and pattern analyses between conditions with different lexical status either at the global level or at the local level to test the dynamics of chunk processing. When comparing the strings that contained global-level lexical chunks (*GwLw, GwLn*) with the strings that did not contain global-level lexical chunk (*GnLw, GnLn*), the significant differences were observed in the electrodes over middle parietal and left frontal-temporal regions (Fig. 4a, indicated by white points) starting around 170 ms (indicated by the red arrow in Fig. 4a). When comparing the strings that contained local-level lexical chunks (*GwLw, GnLw*) with the strings that did not contain local-level lexical chunk (*GwLn, GnLn)*, the significant differences in response amplitudes started much later at the latency around 230 ms in the electrodes over frontal and parietal-occipital regions (indicated by the green arrow in Fig. 4a).

**Figure 4.**
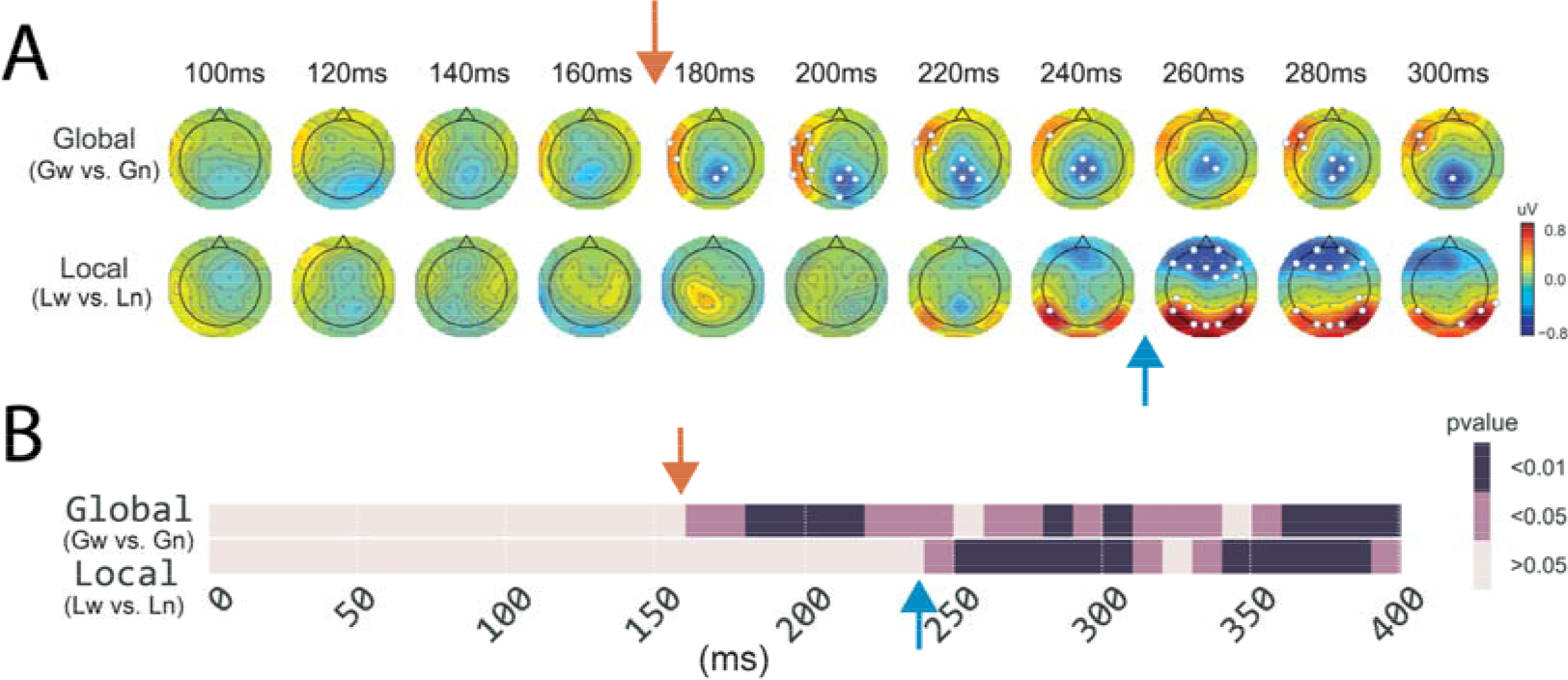
The processing dynamics of chunks at global and local levels. (a) Analysis of response amplitude in topographical comparisons between different lexical status at the Global level (upper row) and at the Local level (lower row) across time. The color scheme depicts the differences in response amplitude between conditions, and the white points superimposed on the topographies indicate the electrodes that showed significant differences after multiple comparison correction (FDR). (b) The temporal dynamics of TANOVA results. The results showed distinct starting time of significant response pattern differences between different lexical status at the Global level and at the Local level. The red arrows in all plots indicate the earliest latency of difference in the Global level comparison, and the green arrows indicate the earliest latency of difference in the Local level comparison.

TANOVA results (Fig. 4b) further showed that response patterns were statistically significantly different between processing the distinct lexical status at the *Global* level began around 170 ms after the stimulus onset (indicated by the red arrow in Fig. 4b), and the significant pattern differences at the *Local* level began around 230 ms (indicated by the green arrow in Fig. 4b), consistent with the observation of significant electrodes in Fig. 4a. All these results suggested that the global-level chunks were processed earlier than chunks at the local-level.

## Discussion

This study investigated the processing dynamics of chunking operation in reading. By using the Chinese four-character strings that contain multiple grain-size language chunks, the behavioral results showed that the recognition of lexicalized local chunks was affected by the lexical status of global chunks, but not vice versa, which suggested that the processing of chunks at the global level was prioritized over the processing of local ones during reading. Moreover, the earliest EEG responses showed distinct patterns between lexicalized and non-lexicalized chunks, and the latency of successive EEG responses was faster when processing chunks at the global level than that for local chunks. These consistent behavioral and electrophysiological results suggested that two distinct stages successively operate in the early stage of reading for detecting and processing chunk information at multiple levels.

### Detection of chunks at 100 ms

In the clustering results (Fig. 2), a ‘temporal gap’ was observed in the early EEG reading responses and separated the processing from 80 to 200 ms into two distinct clusters, suggesting the different neural bases and possible distinct functions. Furthermore, the response patterns of earliest cluster around 100 ms were modulated by the lexical status of chunks at both global and local levels (Fig. 3). These findings are consistent with the early lexical familiarity checking mechanism proposed in the E-Z reader model (Reichle et al., 2003). Language chunks and their lexical status should be checked before accessing the semantics. In other words, the familiar lexical chunks are detected before subsequent processes (e.g. semantic retrieval). This is especially important in the language that lacks explicit boundaries for lexicalized chunks/phrases, such as written Chinese. Our results suggesting such lexical checking/detection can occur early in the reading process around 100ms and extend to multiple chunk levels.

What factor enables this early chunk detection in reading? Top-down mechanisms have been proposed to account for perceptual and cognitive functions, such as the prior knowledge or prediction of the global shape information in object recognition (M. Bar et al., 2006; Moshe Bar, 2003; Panichello, Cheung, & Bar, 2012). The detection of language chunks at multiple levels during reading involved the left frontal regions and occipital regions (Fig. 3a), similar to the top-down modulation by the early feed-forward projection of low spatial frequency information (M. Bar et al., 2006). In previous research, high-frequency words can be easily detected and recognized (N. C. Ellis, 2002; Monsell & Besner, 1991). The transparence (MacGregor & Shtyrov, 2013) and decomposability (Abel, 2003; Vannest, Polk, & Lewis, 2005) also affect the mental encoding of complex words, phrases and idioms. However, individual differences in reading may make the perception of these physical attributes vary in a great degree across individuals. Therefore, the factor that leads to the early chunk detection are likely to be the perceptual consequences -- the familiarity of these attributes. In fact, the familiarity has been demonstrated in improving language retrieval (Bannard & Matthews, 2008; Zheng, Li, & Xiao, 2015). In this study, we controlled the familiarity by only using stimuli that were rated at the extreme degree of familiarity -- either very familiar words or strange nonwords. We speculated that the familiarity of lexical-orthographic features (such as frequency and decomposability) is the criterion of chunk detection, and it can apply simultaneously at both global and local levels during early reading processes.

### Priority of processing global chunks

Our behavioral results demonstrated that the processing of a global chunk significantly affected the lexical decision of lexicalized local chunks. Whereas the local chunks had no impact on the lexical decision of the lexicalized global chunk. The unidirectional effect suggested that the processing of global level chunks had priority over the processing of their constituents. EEG further provided evidence supporting the temporal hierarchy in processing global and local chunks. The EEG results showed that the processing of global chunk started around 170 ms, while the onset of local chunk processing was much later (around 230 ms). These EEG results, together with our behavioral data, demonstrated that after the simultaneous chunk detection at both levels, the processing of different sizes of lexical chunks began at different times: the processing of global chunks preceded that of local chunks.

The priority of global information has been demonstrated in many cognitive domains. Gestaltism (Dewey, n.d.; Heider, 1977) considers the global contains more information than the aggregation of its locals. In vision, the global precedence effect (Navon, 1977/7) suggests that recognizing a scene is hierarchical and global processing has priority over local processing; while local processing is subject to the top-down re-evaluation and integration into global processing. Similarly, the top-down facilitation of visual object recognition also implies the activation of high-level information will be faster than the lower-level information (M. Bar et al., 2006; Moshe Bar, 2003). In linguistics, the word superiority effect (Reicher, 1969) -- advantage of words on recognizing letters -- suggests that the processing of a word at the global level interacts with the letter identification (McClelland & Rumelhart, 1981a). This study further demonstrates the influences of phrases on words. Our results expand previous research and suggest that the global-priority mechanism can be applied across multiple levels in a hierarchical manner in the linguistic context. The priority of global chunk is consistent with the information theory (Shannon 1948): larger chunk contains more context information and less internal entropy, which can prevent ambiguity.

### Paralleled processing of chunks at both levels

The behavioral results revealed that the judgement of a non-lexicalized phrase at the global level was harder when the task-unrelated chunks were familiar words at the local level. This indicated that the local processing may be initiated before the finish of global processing. The EEG results further supported that processing at both levels temporally overlapped -- the response patterns of processing global chunks continued after the start of local processing responses patterns (Fig. 4). This observation of partially temporal overlap in the processing part-whole hierarchies is consistent with simultaneous processing mechanisms implemented in the connectionist networks (Hinton, 1990). A scheduler could control the participation of processing at different levels. Should processing a chunks exceeds expected duration, processing of chunks at other levels would occur. Moreover, the topographic patterns showed left lateralization for processing chunks at the global level, whereas both hemispheres engaged in processing chunks at the local level (Fig. 4), suggesting the possible anatomical differences that mediate the partially temporal paralleled processes at both levels.

Based on all results, we tentatively put forward a workflow of processing multiple-level information in reading (Fig. 5). The segmentation occurs in an early short time window, and possible chunks at all levels are detected based on the familiarity of lexical-orthographic features (detection stage). The chunks at each level are further processed with distinct temporal characteristics (processing stage). Specifically, the processing of global chunks possesses priority over the local chunks, while the processing of local chunks can launch before the finish of global chunk processing. Hence, the processes of chunks at two levels have partially temporal overlap that enables interaction across levels before final integration.

**Figure 5.**
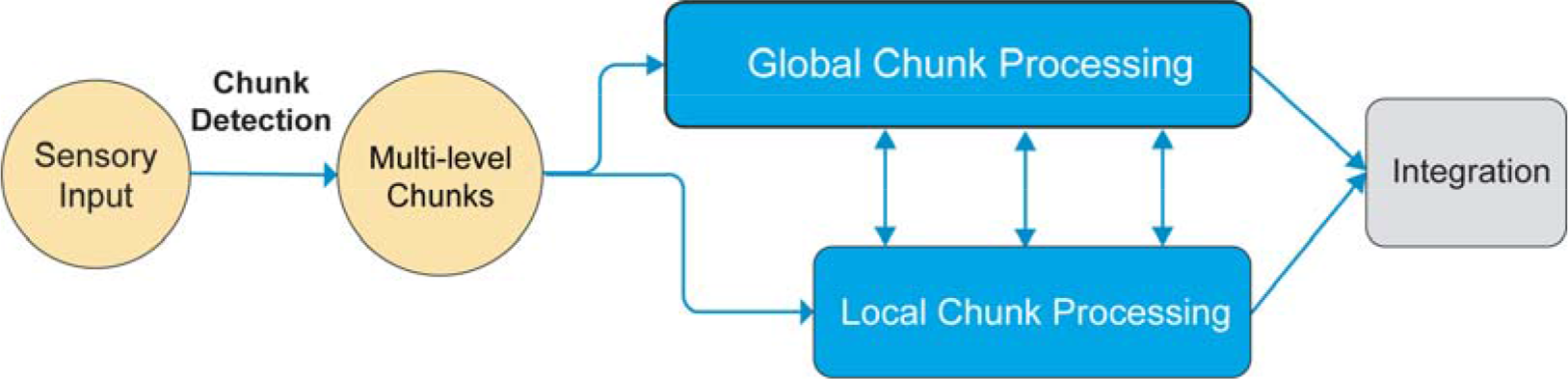
The schematic diagram of proposed two-stage chunking operation in reading.

## Conclusion

The current study investigated the chunking mechanism in reading. Consistent behavioral and EEG results suggested that multiple levels of chunks were realized via two distinct stages of chunking in the early time course of reading. The first stage detected lexicalized chunks at all levels of grain-size, whereas in the second stage the processing of global level led the local level, and resulted later in a parallel and interactive process. This study revealed rich dynamics of chunking operation during reading, which provides the starting computation for comprehension of hierarchical language systems.

## Footnotes

1 The part of speech (POS) is determined by a record of the lexicon of Jieba (Version 0.36; https://github.com/fxsjy/jieba).

2 The frequency was determined by the record of the Sogou Pinyin lexicon, and the high-frequency meant the frequency above 3,000.

3 The character’s log frequency is determined by the Subtitle Database (Cai & Brysbaert, 2010).

